# KK-LC-1 reveals evolutionary divergence in the regulation of sperm motility between humans and mice

**DOI:** 10.64898/2026.07.17.739100

**Authors:** Taiga Yamazaki, Yukiko Yasuoka, Keita Kato, Hiroya Nakamura, Kogiku Shiba, Hiroaki Hamaguchi, Kazuo Inaba, Natsuko Kawano, Tetsuro Yamashita, Takashi Fukuyama

## Abstract

CT83 (KK-LC-1) is a cancer-testis antigen originally identified in human lung cancer cells and has recently attracted attention as a potential target for cancer therapy. Although KK-LC-1 orthologs have been identified in up to 160 animal species, a murine homolog had not previously been identified, hindering *in vivo* analysis of its physiological function. In this study, we identified the mouse homolog of KK-LC-1 and performed a comparative analysis of its properties in humans and mice, together with an investigation of its biological function using gene knockout (KO) mice. The murine *Kk-lc-1* gene is located on the X chromosome and, like its human counterpart, contains an N-terminal transmembrane domain. In both humans and mice, KK-LC-1 is expressed specifically in the testis and localizes to the head and tail regions of sperm.

Analysis of *Kk-lc-1*-deficient mice revealed normal spermatogenesis, and both male and female KO mice were fertile. However, sperm from *Kk-lc-1*-deficient males exhibited reduced motility caused by decreased flexibility of the midpiece and failed to penetrate the oocyte zona pellucida *in vitro*. This defect was rescued by artificial insemination using epididymal sperm, suggesting that maternal factors *in vivo* may compensate for reduced sperm motility. Although impaired sperm motility during *in vitro* fertilization (IVF) was rescued by murine *Kk-lc-1*, functional rescue by human KK-LC-1 was not observed.

These findings indicate that KK-LC-1 contributes to sperm motility but is not essential for fertility. Moreover, species-specific differences in KK-LC-1-mediated regulation of sperm motility suggest functional divergence during evolution. The role of KK-LC-1 in sperm motility should therefore be considered in the clinical development of cancer therapies targeting KK-LC-1.

## Introduction

Cancer-testis antigens (CTAs) comprise a group of genes that are predominantly expressed in testicular cells, including germ cells, and in trophoblasts, while also being aberrantly expressed in a wide range of cancer cells. To date, 277 genes have been registered and reported in the CT database (http://www.cta.lncc.br) [1]. Because of their highly restricted expression in normal tissues and frequent activation in diverse cancers, CTAs are considered attractive targets for cancer therapy [2, 3]. In the development of cancer therapeutics targeting specific molecules, it is essential to elucidate the *in vivo* functions of these molecules and their mechanisms of action in cancer cells. Gene knockout (KO) mouse models play a crucial role in evaluating *in vivo* function and have been instrumental in CTA research. For example, A disintegrin and metalloproteinase 2 (ADAM2), a transmembrane protein that is aberrantly expressed in several malignant tumors, is essential *in vivo* for membrane fusion between sperm and egg during fertilization [4, 5]. Similarly, acrosin-binding protein (ACRBP/OY-TES-1), a CTA expressed in multiple cancer types [6], plays a critical role in acrosome formation during spermatogenesis, and ACRBP deficiency leads to reduced fertility owing to nuclear morphological abnormalities in sperm [7]. These physiological functions have been elucidated through functional analyses of mouse homologs. However, for some clinically important CTAs, such analyses are not feasible. For instance, New York esophageal squamous cell carcinoma-1 (NY-ESO-1) is a highly immunogenic cancer antigen currently under clinical investigation as a therapeutic target [8], yet the absence of a mouse homolog has hindered functional studies using gene KO models.

Kita-Kyushu lung cancer antigen-1 (KK-LC-1), also known as CT83/Cxorf61, is a cancer antigen identified by cytotoxic T lymphocytes (CTLs) from human lung cancer cells [9]. In normal human tissues, KK-LC-1 expression is restricted to the testis, whereas it is aberrantly upregulated in various cancers, including gastric cancer [10] and breast cancer [11]. Recently, KK-LC-1 has attracted increasing attention as a potential target for anticancer therapy [12]. Orthologs of KK-LC-1 have been identified in approximately 160 animal species in the NCBI database, suggesting that KK-LC-1 is a relatively well-conserved protein. Nevertheless, a homologous gene has not been annotated in mice (*Mus musculus*), posing a major obstacle to *in vivo* functional analysis. In this study, we identified *Gm28269* as a candidate murine homolog of KK-LC-1 through *in silico* analyses. Similar to human KK-LC-1, *Gm28269* encodes a protein containing an N-terminal transmembrane domain, is specifically expressed in testicular germ cells, and localizes to both the sperm nucleus and tail. Although *Gm28269* deficiency did not affect fertility under spontaneous mating conditions, it resulted in reduced sperm motility due to impaired flexibility of the sperm midpiece. This defect compromised sperm penetration of the zona pellucida and led to decreased fertilization rates *in vitro*. Notably, this phenotype was rescued by murine *Gm28269* but not by human *KK-LC-1*, indicating species-specific differences in the regulation of sperm motility by KK-LC-1 in humans and mice.

## Results

### Identification and characterization of mouse *Kk-lc-1*

A 27-bp mouse genomic sequence with high homology to the second exon of the human *KK-LC-1* gene was identified using the Multiz Alignment of 100 Vertebrates in the UCSC Genome Browser (Supplementary Fig. 1A, B). Using this sequence as a query, we performed a BLAT search in the Integrative Genomics Viewer (IGV) [13] and identified *Gm28269* as a candidate murine homolog of *KK-LC-1* (Supplementary Fig. 1C). Similar to human *KK-LC-1*, *Gm28269* is located on the X chromosome and consists of two exons arranged in a tail-to-tail orientation adjacent to *SLC6A14* (Fig. 1A).

**Figure 1.**
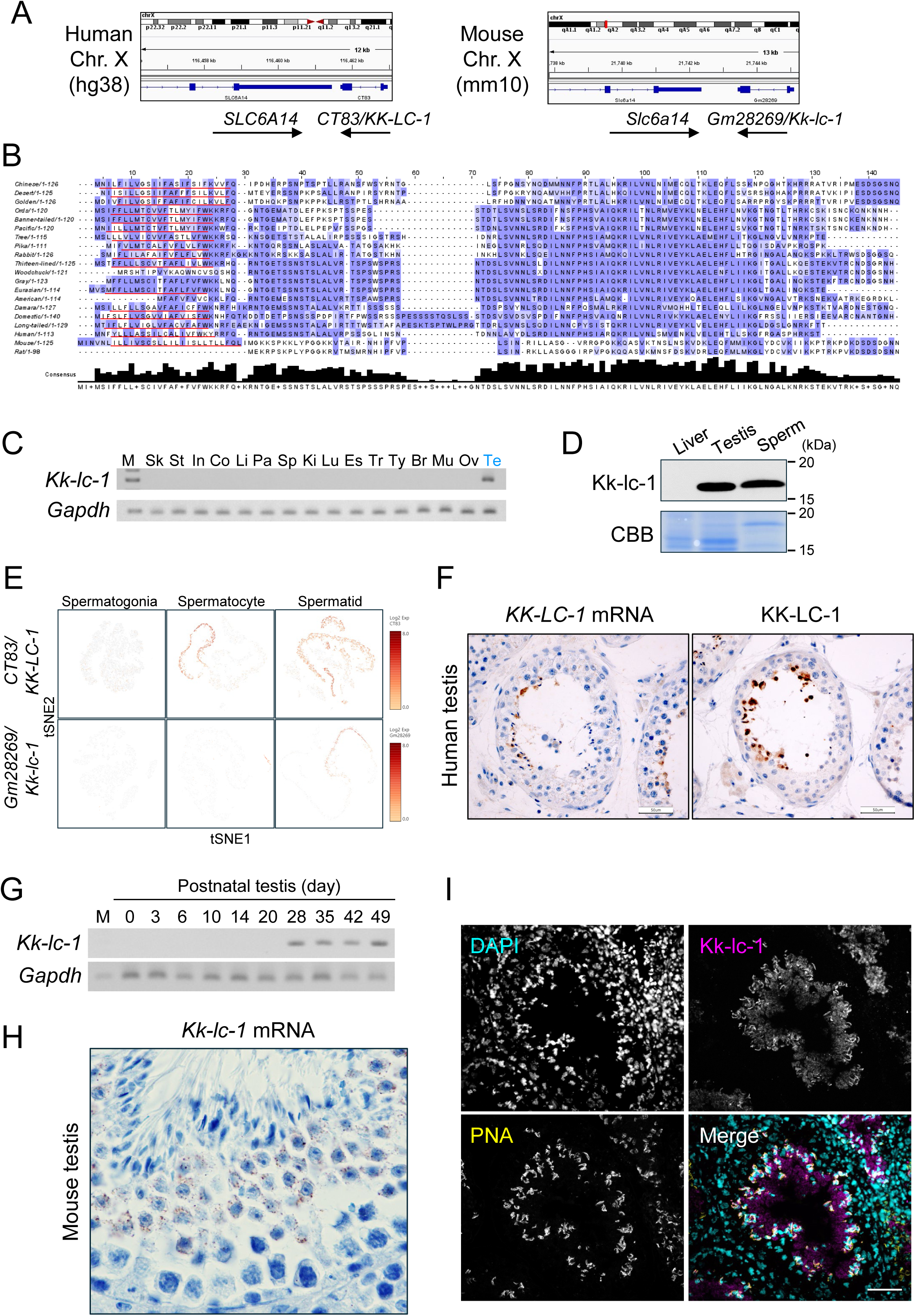
Characterization of mouse homologue of *KK-LC-1*. A. Snapshot of Integrative Genomics Viewer (IGV) from human (hg38) and mouse (mm10) X chromosome. Both *CT83*/*KK-LC-1* and *Gm28269*/*Kk-lc-1* are located on downstream of *SLC14A* gene. B. Amino acid alignment of various species of *CT83*/ *KK-LC-1* and *Gm28269*/*Kk-lc-1* obtained from mouse and rat by CLUSTALW. The amino acids were color-coded according to the Blosum62 matrix. Gaps are colored white. If a residue matches the consensus sequence residue at that position it is colored dark blue. If it does not match the consensus residue but the 2 residues have a positive Blosum62 score, it is colored light blue. Note that transmembrane domain (lined with red bar) was located on most of CT83 and Gm28269 N-terminus. C. Detection of *Kk-lc-1* expression in various mouse tissues by RT-PCR. *Gapdh* expression was detected as control. M: marker, Sk: skin, St: stomach, In: intestine, Co: colon, Li: liver, Pa: pancreas, Sp: spleen, Ki: kidney, Lu: lung, Es: esophagus, Tr: trachea, Ty: thymus, Br: brain, Mu: muscle, Ov: ovary, Te: testis. D. Detection of Kk-lc-1 by immunoblot. E. Stage-specific expression of *KK-LC-1* in the human and mouse testis. ScRNA-seq data deposited by Hermann et al. were reanalyzed. The t-SNE plot shows the distribution of cells expressing the *KK-LC-1* at each developmental stage of testicular germ cells. F. Detection of KK-LC-1 expression in human testis. Left panel shows transcripts of KK-LC-1 detected by *in situ* HCR. Right panel shows immunostaining of KK-LC-1. Scale bar, 50 μm. G. Detection of *Kk-lc-1* expression in newborn mouse testis. *Kk-lc-1* expression was detected by RT-PCR. *Gapdh* expression was detected as control. M: marker. H. Detection of *Kk-lc-1* mRNA in adult mouse testis by *in situ* Hybridization chain reaction (HCR). *Kk-lc-1* mRNA was observed as particle signal in testicular cells. Scale bar, 20 μm. I. Expression of *Kk-lc-1* protein in adult mouse testis. A snapshot of the seminiferous tubule was shown. Kk-lc-1 was detected by anti-Kk-lc-1 antibody (pink) and acrosome and DNA were stained with PNA (yellow) and DAPI (cyan), respectively. Scale bar, 50 μm.

Several rodent species have been reported to possess *KK-LC-1*, including members of Castorimorpha (e.g., American beaver, North American beaver, banner-tailed kangaroo rat), Sciuridae (e.g., Eurasian red squirrel, gray squirrel, thirteen-lined ground squirrel, woodchuck), Hystricognathi (e.g., Damara porcupine, crested porcupine, long-tailed porcupine), and Cricetidae (e.g., Chinese hamster, desert hamster, golden hamster). We compared the amino acid sequences of KK-LC-1 from these species with mouse and rat Gm28269, as well as with KK-LC-1 from Scandentia (tree shrews), Lagomorpha (pikas and rabbits), and Primates (*Homo sapiens*) (Fig. 1B).

Sequence alignment revealed that mouse and rat Gm28269 share relatively high sequence similarity, whereas mouse Gm28269 and human KK-LC-1 share approximately 20% amino acid identity, with limited conservation of specific sequence motifs. Phylogenetic analysis further indicated that Gm28269 in Muridae (mouse and rat) is more evolutionarily divergent than KK-LC-1 in other rodent lineages, while remaining relatively conserved within the *Mus* genus (Supplementary Fig. 2A, B). Despite the low overall sequence homology between human KK-LC-1 and mouse Gm28269, both proteins possess an N-terminal transmembrane domain (Fig. 1B). Consistent with this structural similarity, KK-LC-1 is a small protein with a molecular weight of 12.8 kDa, and Gm28269 likewise has a low molecular weight of 13.8 kDa.

We next examined the tissue expression profile of *Gm28269* by RT-PCR and found that its expression is restricted to the testis (Fig. 1C). At the protein level, Gm28269 was detected in both the testis and sperm (Fig. 1D). Although testis-specific expression of KK-LC-1 has been reported previously, the precise stage of germ cell differentiation at which expression occurs has remained unclear. To address this issue, we reanalyzed published intratesticular single-cell RNA sequencing (scRNA-seq) datasets to assess the expression of *KK-LC-1* and *Gm28269* across spermatogonia, spermatocytes, and spermatids (Fig. 1E). In humans, a distinct cell population exhibited robust upregulation of *KK-LC-1* beginning at the spermatocyte stage, with expression detected in the majority of spermatids. In mice, *Gm28269* expression was observed in only a small subset of spermatocytes but was detected in more than half of spermatids.

To further validate these findings, we analyzed human testis tissue panels at both the transcript and protein levels. KK-LC-1 expression and localization were confirmed in spermatocytes and spermatids (Fig. 1F). In mice, RT-PCR analysis of neonatal testis samples revealed that *Gm28269* expression initiates at postnatal day 28 (Fig. 1G), corresponding to the appearance of haploid cells during the first wave of spermatogenesis, consistent with the scRNA-seq results. Moreover, RNA in situ hybridization in testis sections confirmed *Gm28269* expression in spermatids, where punctate nuclear signals were observed in round spermatids identified by hematoxylin counterstaining (Fig. 1H). At the protein level, Gm28269 localized to the sperm head, distinct from the acrosome, and was also detected in the lumen of seminiferous tubules (Fig. 1I).

Collectively, based on its chromosomal location, conserved N-terminal transmembrane domain, testis-specific expression, and spermatid-enriched localization, we conclude that Gm28269 represents the murine homolog of human KK-LC-1.

### KK-LC-1 localizes both sperm head and sperm tail

To characterize the subcellular localization of KK-LC-1 in sperm, we performed biochemical fractionation analyses using human and mouse spermatozoa. KK-LC-1 was not solubilized by RIPA buffer containing 1% Triton X-100 and 0.1% SDS in either human or mouse sperm (Fig. 2A), indicating that KK-LC-1 is not associated with the sperm plasma membrane. Instead, it is likely enriched in insoluble structures, such as the sperm flagella [14] or nucleus-associated domains [15]. To further define its localization, spermatozoa were fractionated into head and tail components (Fig. 2B, C), and whole lysates from each fraction were analyzed by immunoblotting. KK-LC-1 was detected in both the head and tail fractions of human and mouse spermatozoa (Fig. 2D). To determine its precise localization within the sperm head, mouse spermatozoa were further fractionated into the apical region, perinuclear theca (PT), and sperm nuclear fractions (Fig. 2E). The distribution of Kk-lc-1 among these fractions was examined by immunoblotting. Successful fractionation was verified using ADAM3 as a marker for the sperm membrane and ACTRT2 as a marker for the PT, confirming effective separation of the subcellular components. Kk-lc-1 was predominantly detected in the sperm nuclear fraction, which represents a more insoluble component of the sperm head compared with the PT fraction (Fig. 2F).

**Figure 2.**
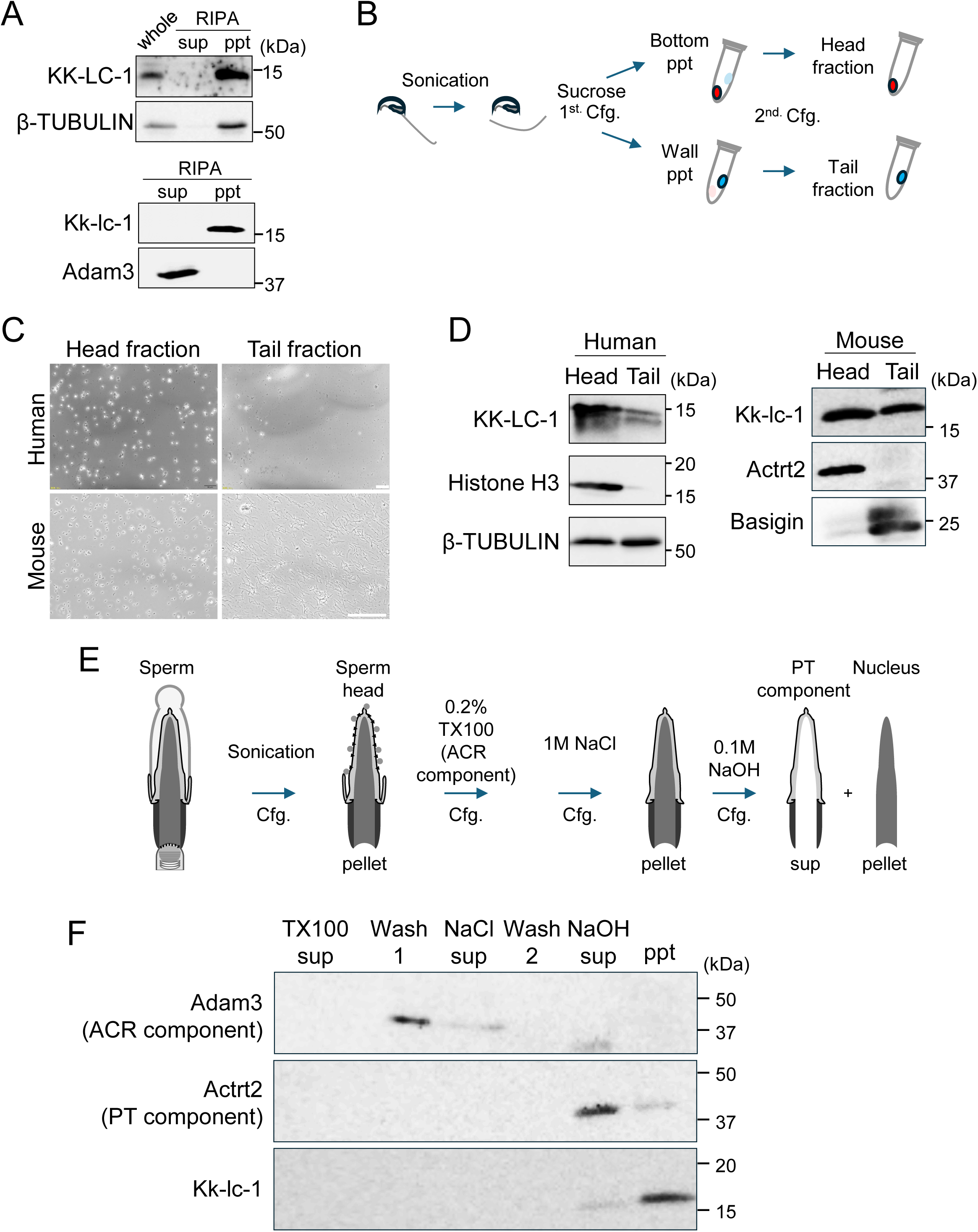
KK-LC-1 localizes in the sperm nucleus and the sperm tail. A. RIPA extraction of sperm KK-LC-1. Human (upper panel) and mouse (lower panel) sperm were extracted with RIPA buffer. Both the RIPA supernatant (sup) and pellet (ppt) fractions were subjected to immunoblotting. Whole lysate (whole) was also subjected in human sample. B. Scheme of sperm head/tail fractionation. Sonicated sperm were fractionated by sucrose buffer followed by two steps of centrifugation. C. Snapshot of sperm head and tail fractions. Scale bar 50 μm (human) and 80 μm (mouse). D. Localization of KK-LC-1 in sperm head and tail. Validation of fractionation was performed with detection of sperm head protein histone H3 (for human sperm), Actrt2 (for mouse sperm) and sperm tail protein β-TUBLIN (for human sperm), Basigin (for mouse sperm). Note that KK-LC-1 was detected in both head and tail fractions of sperm in human and mouse. E. Fractionation of mouse sperm head proteins. Serial fractionation method reported by Zhang et al. enables to prepare acrosomal proteins (ACR) and proteins contained with perinuclear theca (PT). F. Localization of Kk-lc-1protein in mouse sperm head. Fractionation was validated with Adam3 (ACR protein) and Actrt2 (PT protein). Note that most Kk-lc-1 localizes NaOH pellet (ppt) after PT extraction.

### The reduced IVF rate caused by Kk-lc-1 deficiency is rescued by maternal environment

To investigate the *in vivo* function of *Gm28269/Kk-lc-1*, we generated *Kk-lc-1* knockout (KO) mice using CRISPR/Cas9-mediated genome editing (Fig. 3A). Frameshift mutations in *Kk-lc-1* were confirmed by genomic analysis (Fig. 3B), and loss of Kk-lc-1 expression was verified by immunoblotting (Fig. 3C) and immunostaining of testis sections (Fig. 3D). No significant differences were observed in testis morphology or weight between *Kk-lc-1*-deficient and wild-type (WT) male mice (Fig. 3E). Furthermore, fertility assessment by natural mating revealed comparable litter sizes between WT and KO mice (Fig. 3F).

**Figure 3.**
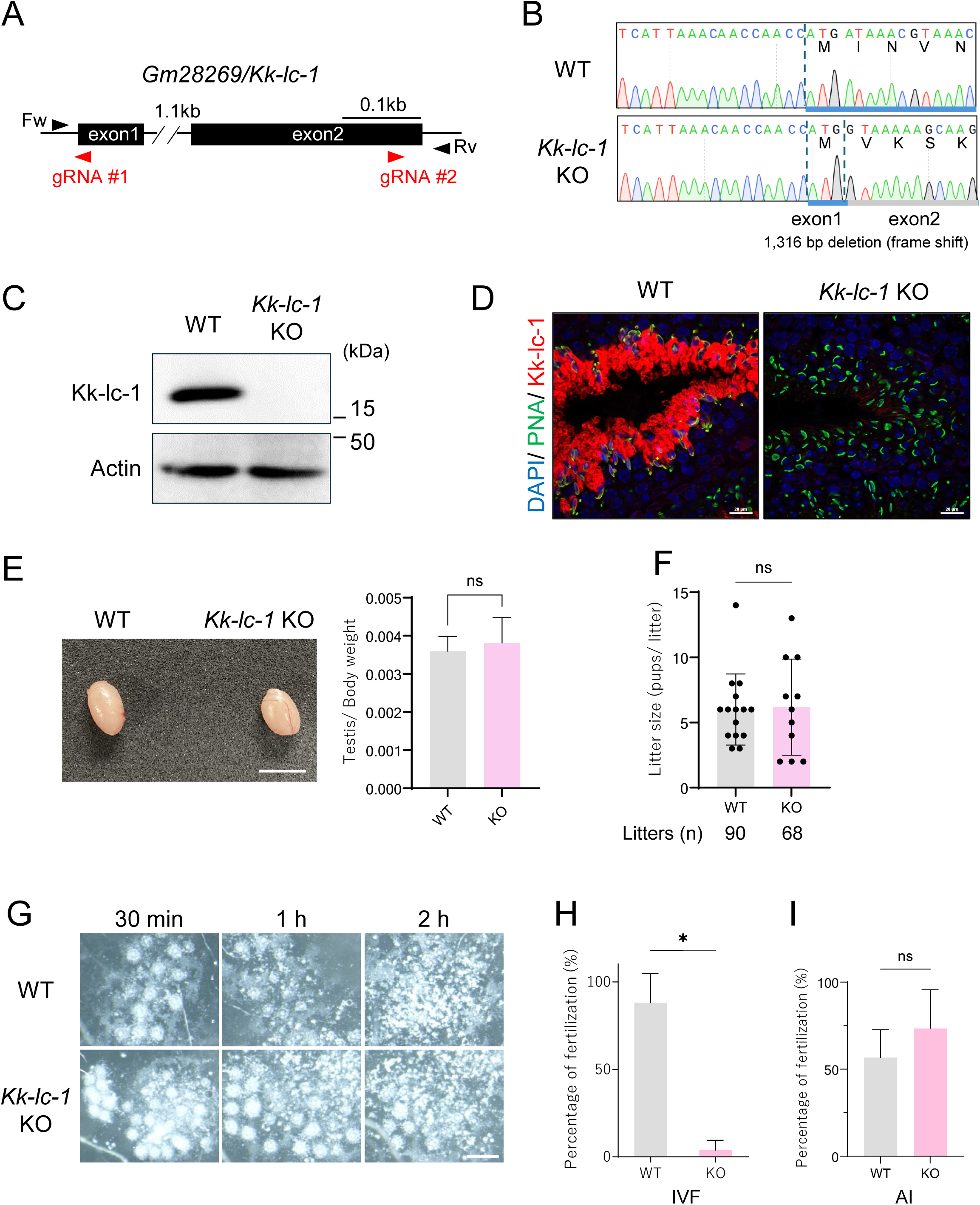
The fertility of *Gm28269*/*Kk-lc-1* deficient male mice is influenced by maternal environment. A. CRISPR/Cas9 targeting scheme. Two gRNAs (red arrowheads) were designed to target exons 1 and 2. Fw and Rv (black arrowheads) are forward and reverse primers for genotyping, amplicon sequencing. B. Indel sequence result of the *Kk-lc-1* knockout allele. Large deletion (1,316 bp) and frame shift occurred in *Kk-lc-1* knockout allele. C. Detection of Kk-lc-1 in WT and *Kk-lc-1* KO testis by immunoblotting. Actin was used as a loading control. D. Detection of Kk-lc-1 in WT and *Kk-lc-1* KO testis by immunostaining. PNA (green) and DAPI (blue) were used to visualize the acrosome and DNA, respectively. Scale bar, 20 μm. E. Testicular morphology and weight of *Kk-lc-1* KO mice testes. Ratio of Testis/body weight were presented. Unpaired *t*-test reveals no significant difference (ns) between WT and *Kk-lc-1* KO testis weight. Scale bar, 4 mm. F. Number of litter size per pregnancy was presented. Two males each for WT and *Kk-lc-1* KO were mated with WT female. No significant difference (ns) was observed between WT and *Kk-lc-1* KO litter size by unpaired *t*-test. G. Cumulus cell layer dispersion after sperm addition to cumulus-oocyte complex (COCs). Snapshots of COCs at 30 minutes, 1 hour, 2 hours after insemination were presented. Scale bar, 0.5 mm. H. Fertility of *in vitro* fertilization (IVF) with *Kk-lc-1* KO sperm. Cumulus-intact oocytes were inseminated with WT or *Kk-lc-1* KO sperm. The fertility rate was calculated based on 2-cell development rate \**P* < 0.05 (unpaired *t*-test). I. Fertility of artificial insemination with *Kk-lc-1* KO sperm. Cauda epididymal sperm were injected into uterus. The fertility rate was calculated based on 2-cell development rate. ns: not significant (unpaired *t*-test).

In contrast, during *in vitro* fertilization (IVF) assays using *Kk-lc-1*-deficient sperm, dispersion of cumulus cells was delayed following sperm addition compared with WT sperm (Fig. 3G), and the fertilization rate was significantly reduced (Fig. 3H). These observations suggest that *in vivo* reproductive conditions partially compensate for the fertilization defects associated with *Kk-lc-1* deficiency.

To determine whether this compensatory effect originates from the male reproductive tract (e.g., seminal plasma components) or from the female reproductive environment (e.g., uterine or oviductal factors), epididymal sperm from *Kk-lc-1*-deficient males—thus lacking seminal plasma—were artificially inseminated into the female uterus. Notably, the reduced fertilization rate observed in IVF was restored by artificial insemination (Fig. 3I), indicating that maternal factors are sufficient to compensate for the fertilization defects of *Kk-lc-1*-deficient sperm during *in vivo* fertilization.

### *Kk-lc-1*-deficient sperm exhibit impaired zona pellucida penetration associated with reduced midpiece flexibility *in vitro*

Given that *Kk-lc-1* localizes to both the sperm nucleus and tail, we hypothesized that its loss might affect sperm morphology. To test this, we compared the morphology of wild-type (WT) and *Kk-lc-1* knockout (KO) spermatozoa by fluorescence staining of mitochondria and nuclei. No obvious morphological abnormalities were detected, and KO sperm exhibited grossly normal morphology (Fig. 4A). We next evaluated sperm motility using computer-assisted sperm analysis (CASA). Motility parameters, including average path velocity (VAP), curvilinear velocity (VCL), and straight-line velocity (VSL), were measured after 1 h of incubation. *Kk-lc-1*-deficient sperm showed significantly reduced motility across all parameters compared with WT sperm (Fig. 4B). In addition, the proportion of sperm exhibiting hyperactivated motility was significantly lower in KO sperm than in WT sperm from the onset of incubation through 2 h post-incubation (Fig. 4C). High-speed video microscopy revealed that *Kk-lc-1*-deficient sperm displayed reduced flexibility of the flagellum in the region proximal to the sperm head (Fig. 4D, E). Quantitative analysis confirmed a significant decrease in the amplitude of flagellar curvature in KO sperm (Fig. 4F).

**Figure 4.**
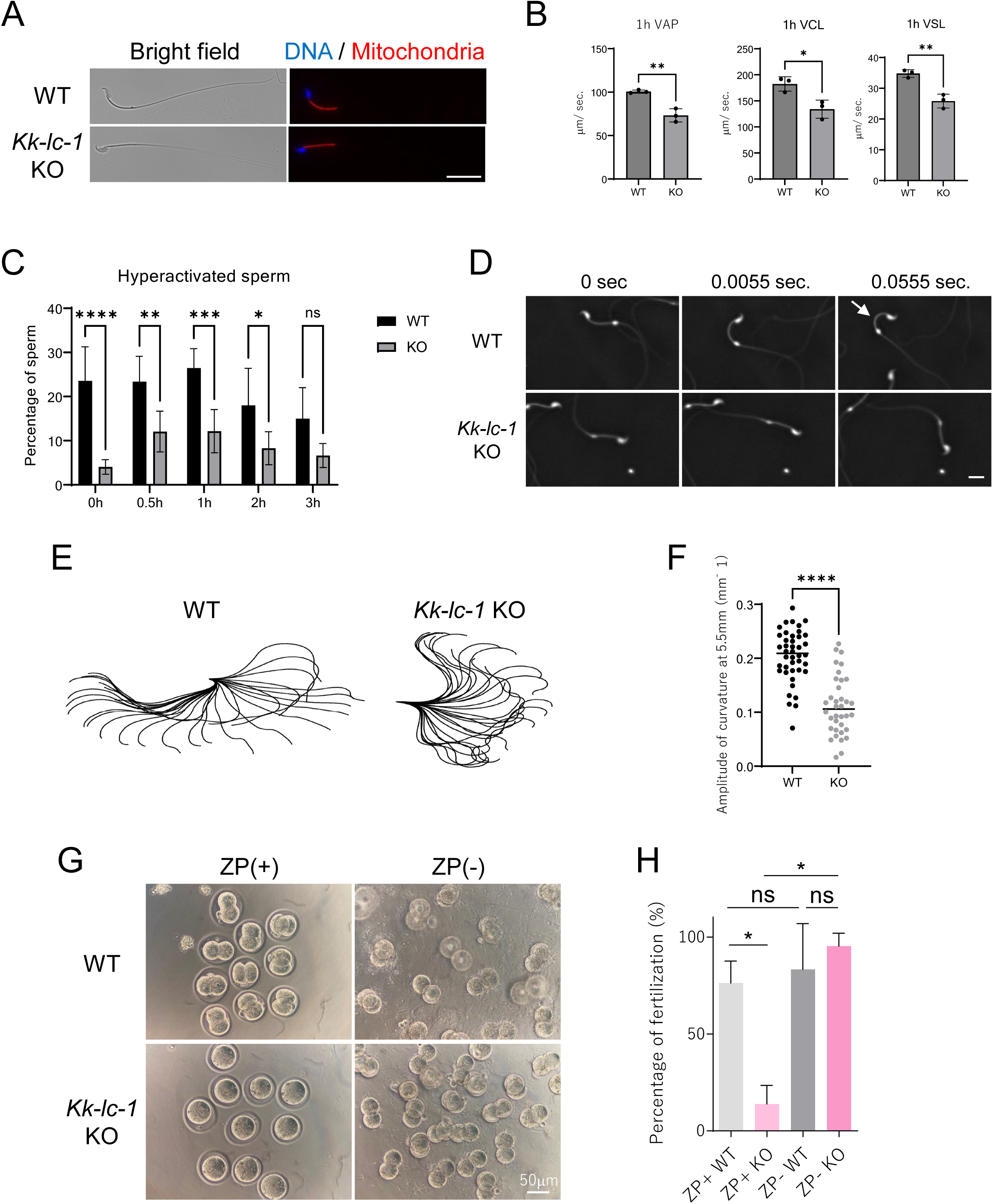
*Gm28269*/*Kk-lc-1* deficient spermatozoa have impaired motility and ZP penetration *in vitro*. A. Morphology of *Gm28269*/*Kk-lc-1* KO sperm. Cauda epididymal sperm were obtained and stained with DNA (Hoechst 33342, blue) and Mitochondria (MitoBright LT Red, red). Scale bar, 20 μm. B. Quantification of sperm motility using SMAS. VAP, VCL and VSL of sperm 1 hour after incubation was plotted \**P* < 0.05, \*\**P* < 0.01 (unpaired *t*-test). C. Temporal changes in the abundance of hyperactivated sperm. At each time point, spermatozoa were analyzed by SMAS and resulted scores were calculated to judge the sperm were hyperactivated or not. \**P* < 0.05, \*\**P* < 0.01, \*\*\**P*<0.001, \*\*\*\**P*<0.0001, ns: not significant (Two-way ANOVA). D. Time-lapse snapshots of individual sperm 1 hour after incubation. The arrow indicates the regions where bending was observed in the sperm tail. Scale bar, 10 μm. E. Sperm flagellar bending patterns are superimposed for 0.166 seconds (30 frames). The origin of the coordinates shows the sperm head base. F. Quantification of the amplitude of curvature of sperm tail. The amplitude of flagellar curvature at the position 5.5 μm from the sperm head base, where frequent tail bending was observed were plotted. \*\*\*\**P*<0.0001 (unpaired *t*-test). G. Snapshots of ZP-free embryos 20 hours after insemination. ZP-intact or -removed oocytes were inseminated with WT or *Gm28269*/*Kk-lc-1* deficient sperm *in vitro*. Scale bar, 50 μm. H. Fertilization rate of ZP-free oocytes. The development rate to the 2-cell stage was used as the fertilization rate. \**P* < 0.05, ns: not significant (unpaired *t*-test).

Sperm motility, including flagellar flexibility and hyperactivation, is crucial for sperm to penetrate the oocyte zona pellucida, in addition to undergoing the acrosome reaction (AR). We next examined whether *Kk-lc-1*-deficient sperm retained fertilization capacity when zona penetration was bypassed. To this end, we performed *in vitro* fertilization (IVF) using zona-free oocytes (Fig. 4G, H). Under these conditions, the fertilization rate of *Kk-lc-1*-deficient sperm exceeded 90%, comparable to that of WT sperm. These results indicate that the reduced fertilization efficiency observed in IVF with intact oocytes is primarily attributable to impaired zona pellucida penetration caused by defective sperm motility in the absence of *Kk-lc-1*.

### Verification of functional rescue of Kk-lc-1 by human KK-LC-1

To confirm whether the impaired sperm motility and reduced IVF success rate observed in *Kk-lc-1*-deficient mice were directly caused by *Kk-lc-1* deficiency, we generated *Kk-lc-1* knock-in (KI) mice using CRISPR/Cas9 (Fig. 5A). Additionally, to investigate whether the sperm dysfunction caused by *Kk-lc-1* deficiency could be rescued by human *KK-LC-1*, we simultaneously generated KI mice expressing human *KK-LC-1* at the *Kk-lc-1*-deficient allele on X chromosome (Fig. 5A). RT-PCR analysis confirmed that the testes of the generated mice (Fig. 5B) expressed transcripts derived from the knock-in gene at the *Kk-lc-1* KO locus (Fig. 5C). Furthermore, protein expressions were also detected in those *Kk-lc-1* and *KK-LC-1* KI testes (Fig. 5D).

**Figure 5.**
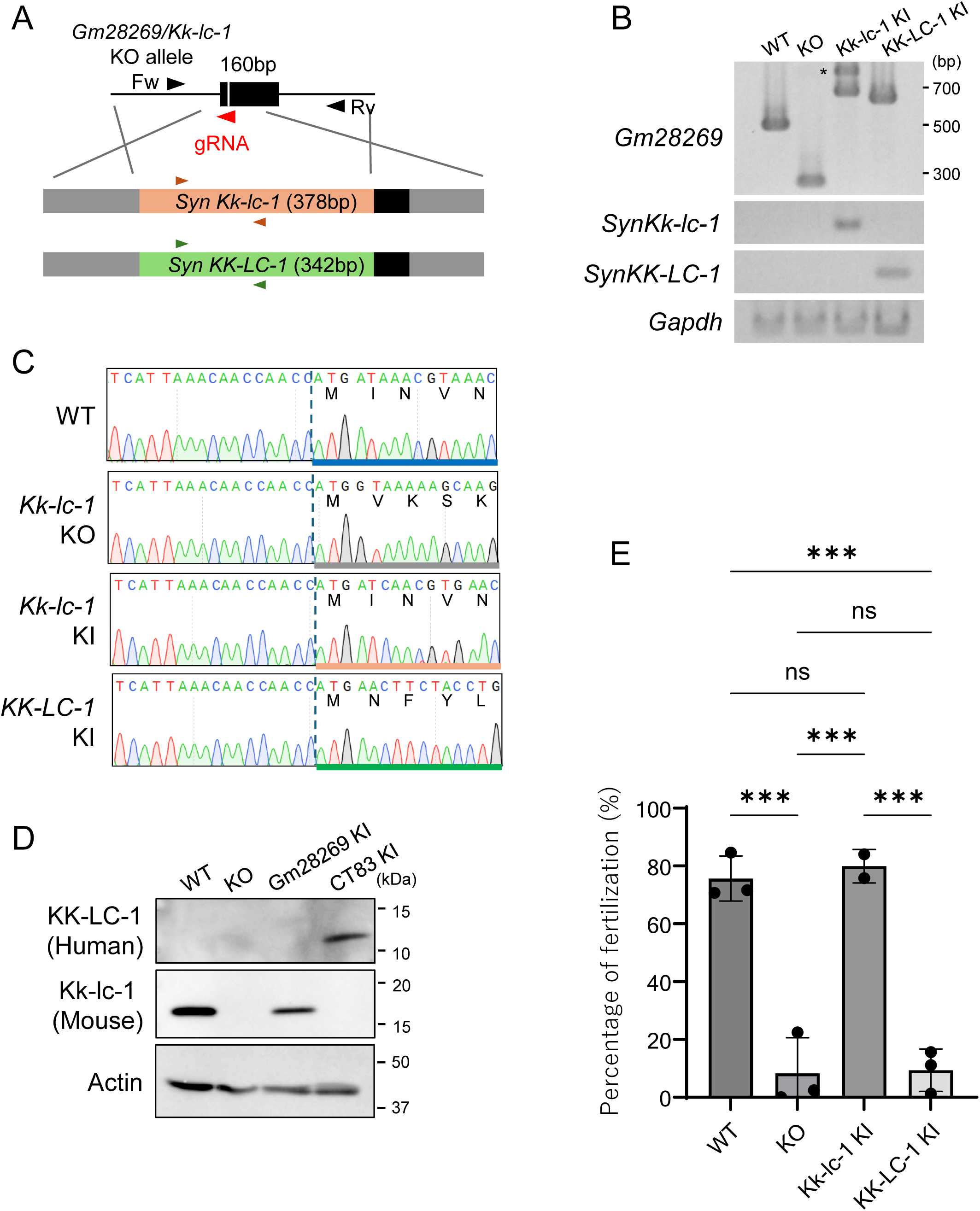
Verification of functional rescue of mouse *Kk-lc-1* by human *KK-LC-1*. A. CRISPR/Cas9 targeting scheme for producing *Gm28269*/*Kk-kc-1* knock-in and *CT83*/*KK-KC-1* knock-in mice. A gRNA (red arrowhead) was designed to target *Kk-lc-1* KO allele and ssODN were used for knock-in DNA. Fw and Rv (black arrowheads) are forward and reverse primers for genotyping, amplicon sequencing. Brown and green arrowheads represent codon optimized synthetic *Kk-lc-1* and *KK-LC-1* specific primer sets, respectively. B. Detection of transcripts from *Kk-lc-1* allele. RT-PCR was performed with testicular RNA. *Gapdh* expression was detected as control. The asterisk indicates non-specific amplification product. C. Sequencing result of *Kk-lc-1* allele. Large deletion (1,316 bp) and frame shift occurred in *Kk-lc-1* knockout allele. E. Fertilization rate of *Kk-lc-1* and *KK-LC-1* KI mice. The development rate to the 2-cell stage was used as the fertilization rate. \*\*\**P* < 0.001, ns: not significant (One-way ANOVA).

Evaluation of male reproductive capability showed that both *Kk-lc-1* KI and *KK-LC-1* KI male mice displayed normal fertility, producing typical litter sizes through natural mating. To assess whether the knock-in genes could restore the fertilization ability of KO sperm, we carried out IVF assays. The results demonstrated that the low fertilization rate observed in *Kk-lc-1* KO mice was fully rescued by *Kk-lc-1* KI, restoring the fertilization rate to levels comparable to WT mice (Fig. 5E). In contrast, *KK-LC-1* KI failed to rescue the low fertilization rate in *Kk-lc-1* KO mice, with the fertilization rate remaining at approximately 20%, similar to that of *Kk-lc-1* KO mice (Fig. 5E).

## Discussion

In this study, we identified *Gm28269* as a potential mouse homolog of *KK-LC-1* through *in silico* analysis based on the human *KK-LC-1* DNA sequence. Phylogenetic analysis of rodents, based on amino acid sequences, revealed that the amino acid sequences of Gm28269 and KK-LC-1 are relatively well conserved within each taxonomic group, including Castorimorpha, Sciuridae, Hystricomorpha, Cricetidae, and Muridae (Fig. 1B and Supplemental Fig. 2A). Additionally, the amino acid sequence of Kk-lc-1 in the genus *Mus* was found to be relatively distinct from those of other rodent families and suborders (Supplemental Fig. 2A). Molecular phylogenetic analysis of *Rodentia* conducted by Huchon et al. revealed that the suborder *Castorimorpha* diverged from *Myodonta* and *Anomaluromorpha* approximately 67.8 million years ago during the Cretaceous period [13]. This suggests that genetic sequence changes in *Kk-lc-1* and *KK-LC-1* may have occurred as early as this time. Notably, the presence of an Lx6 sequence, belonging to the L1 family (data not shown), within an intron of gene A suggests that this gene may have been influenced by retrotransposons during the course of evolution.

This study revealed that Kk-lc-1 plays a functional role in sperm motility but does not affect fertility *in vivo*. In mice, several molecules involved in maintaining the flexibility of the sperm midpiece and regulating motility have been reported, including the sperm-specific calcineurin subunits PPP3CC and PPP3R2 [16], as well as SPATA33 [17]. Notably, deficiencies in any of these molecules result in reduced fertility. On the other hand, Kk-lc-1 is likely to regulate the flexibility of the sperm midpiece through a distinct mechanism from these molecules. Additionally, glycosylation of microtubules in the sperm flagellum has been reported to play a role in flagellar motility [18]. In double-KO mice lacking the tubulin-tyrosine ligase-like (TTLL) family members TTLL3 and TTLL8, tubulin glycosylation in the sperm tail is completely absent. These mice exhibit a subfertile phenotype, similar to Kk-lc-1 KO mice phenotype, characterized by a slight reduction in fertility rather than complete infertility, along with decreased *in vitro* fertilization rates and abnormal bending of the sperm midpiece [18].

The phenotype of Kk-lc-1 KO mice includes a low fertilization rate in *in vitro* fertilization, whereas no fertility issues are observed in natural mating. When sperm collected from the cauda epididymis of KO mice is used for intrauterine insemination, the fertilization rate is comparable to that of WT mice, suggesting the presence of a maternal environment that enhances fertilization. The role of the maternal environment in promoting fertilization has also been suggested by analyses of Prss21-deficient mice [19] and Prss21/Acrosin double-knockout mice [20], as well as by the importance of oviduct-secreted OVGP1 in fertilization and early development in hamsters [21]. Although Kk-lc-1 deficiency impairs sperm motility, the mechanism by which maternal factors enhance the fertilization rate of Kk-lc-1-deficient sperm remains unknown and requires further investigation.

The inability of the human gene to rescue the low fertilization rate observed during *in vitro* fertilization using KO mice suggests the presence of species-specific differences in the regulation of sperm flagellar motility [22]. The initiation of sperm flagellar movement primarily depends on an increase in intracellular pH and the influx of calcium ions. However, human sperm exhibit higher proton currents (*Hv* currents) mediated by *Hv* channels compared to mouse sperm [23]. This suggests that the mechanisms governing intracellular pH regulation, which are critical for motility initiation, may differ between species. Another possibility is that genes with low amino acid homology between humans and mice may not have conserved functions across species, making it less likely for a phenotype to appear upon gene deletion. The homology of CTAs between mice and humans is low. In particular, for the human CTA, NY-ESO-1, a mouse homolog has not even been identified. It has been reported that the amino acid sequence of the human SSX3 gene shares less than 35% similarity with the mouse Ssxa1, Ssxb1, and Ssxb2 genes [24]. Also, the cancer-testis antigen Mage-A3, located on the X chromosome, shares only about 20% homology between humans and mice [25]. Moreover, there are relatively few examples in which gene deficiency affects reproduction. Mage-A3 belongs to the Mage-A gene cluster on the X chromosome. Interestingly, mice in which all six Mage-A family genes including *Mage-A1*, *Mage-A2*, *Mage-A3*, *Mage-A5*, *Mage-A6*, and *Mage-A8* within this ∼210 kb cluster were deleted did not exhibit infertility, although a slight increase in apoptotic cells within the testes was observed [26]. Additionally, conditional knockout analysis of PRAME, another cancer-testis antigen on the X chromosome, revealed a reduction in the number of germ cells in the testes [27]. However, this did not affect litter size. Even among rodents, certain phenomena may only become apparent when using genome-edited animal models [28] of species more closely related to humans than mice, such as hamsters. For example, acrosin is a protease abundantly present in the sperm acrosome. While acrosin knockout (KO) mice do not exhibit infertility [29], acrosin KO golden hamsters are unable to penetrate the zona pellucida and are infertile [30]. Similarly, knockout of oviduct-specific glycoprotein-1 (OVGP1), a secretory glycoprotein specific to the oviduct, does not reduce fertility in female mice [31]. However, in female OVGP1 KO golden hamsters, normal embryo development and implantation fail, leading to infertility [21]. These findings suggest that studies using animal species more closely related to humans could provide novel insights into KK-LC-1.

Since anti–KK-LC-1 activity has been reported to be useful as a prognostic indicator in clinical settings [32], the development of therapeutic agents targeting KK-LC-1 has been actively pursued [12, 33, 34]. The aim of our investigation into mouse *Kk-lc-1* was to establish an experimental platform for evaluating the testicular toxicity of KK-LC-1–targeted therapeutics using mice. KK-LC-1 has recently attracted considerable attention as a target for both immunotherapies and small-molecule–based treatments. However, as demonstrated in the present study, the amino acid sequence homology between mouse Kk-lc-1 and human KK-LC-1 is markedly low. This finding indicates that assessing testicular toxicity of KK-LC-1–targeted therapies using conventional mouse models is inherently challenging. The human *KK-LC-1* knock-in mice generated in this study overcome this limitation and therefore represent a valuable bioresource for preliminary toxicity evaluation during the development of KK-LC-1–targeted therapeutics. Consequently, this model is expected to facilitate and promote the development of diverse therapeutic strategies targeting KK-LC-1.

## Materials and methods

### Animals

All animal experiments were conducted with the approval of the Animal Experiment Committee at Kitasato University, Iwate University and Meiji University. Mice were purchased from Oriental Yeast Co., Ltd. or Japan SLC, Inc.

### Human samples and ethics

All experiments involving human samples were reviewed and approved by the ethics committee of Kitasato University Medical Center (Approval number: 2024009) and Iwate University (Approval number: 202435) in accordance with ethical guidelines for the handling of human cells. The human testis blocks were purchased from TissueArray.Com LLC.

### Identification of mouse homolog of *CT83*/*KK-LC-1*

From the ‘Multiz Alignments of 100 Vertebrates’ on the UCSC Genome Browser (hg38) for *CT83*/*KK-LC-1*, a 27-base-long sequence “GGTGATTTTTTACCCATGATCAACTGT” was obtained from the exon 2 of *CT83*/*KK-LC-1*, which appeared to have high homology with the mouse genome. Using this sequence information, homologous sequences in the mouse genome (GRCm38/mm10) were searched using the BLAT function in IGV (Broad Institute) [13], leading to the identification of an uncharacterized gene matching the 27-base sequence.

### Sperm head-tail separation

Sperm head-tail separation was performed as previously described [35, 36] with minor modifications. Briefly, human spermatozoa were suspended in 200 µL of phosphate buffer saline (PBS) and subjected to sonication for 10 seconds, repeated four times. Following sonication, 800 µL of a 100% sucrose aqueous solution was added to the sperm suspension, and the mixture was subjected to ultracentrifugation at 200,000 × g for 1 hour under 4 ℃. After ultracentrifugation, the pellet fraction at the bottom outside of the tube was resuspended in PBS and designated as the head fraction. Similarly, the pellet fraction adhered to the inner wall of the tube was resuspended in PBS and designated as the tail fraction. Both the head and tail fractions were centrifuged again at 20,400 × g (15,000 rpm) for 20 minutes under 4 ℃. The head and tail fractions were then collected from the bottom of the tube and processed as head and tail fraction samples.

For mouse sperm, spermatozoa collected from eighteen mice were suspended in 450 µL of PBS and subjected to sonication for 10 seconds. The sonicated sperm suspension was centrifuged at 20,400 × g (15,000 rpm) for 5 minutes under 4 ℃, and the resulting pellet was resuspended in an 80% sucrose solution. The subsequent fractionation of the head and tail fractions was performed in the same manner as for the human samples. After ultracentrifugation, obtained head and tail fractions were centrifuged at 20,400 × g (15,000 rpm) 20 minutes under 4 ℃. The resulting head and tail pellets were resuspended separately in an 80% sucrose solution and subjected to ultracentrifugation at 200,000 × g for 1 hour under 4 ℃. After ultracentrifugation, head and tail collected from tubes in which head and tail were suspended, respectively, in the same manner as before. Half of the head suspension was used for following fractionation of sperm ACR and PT components. Another half of head suspension and tail suspension were centrifuged at 20,400 × g (15,000 rpm) 20 minutes under 4 ℃. The head and tail fractions were then collected from the bottom of the tube and processed as head and tail fraction samples.

### Fractionation of sperm ACR and PT components

Fractionation of sperm ACR and PT proteins was performed according to the method reported by Ferrer et al. and Zhang et al. [37, 15]. Briefly, cauda epidydimal sperm were recovered from ICR male mice (7-12 weeks), the fractionated sperm heads collected from 9 mice were suspended in PBS. The resulting pellet was resuspended in 50 μL of 0.2% Triton X-100 and incubated at 4°C for 1 hour on a rotator. After incubation, the suspension was centrifuged at 2,500 × g for 10 minutes, and the supernatant was collected as the Triton X-100 fraction (TX100 sup). The pellet was washed with 50 μL of PBS, and the supernatant obtained was designated as the Wash 1 fraction. The pellet was then resuspended in 50 μL of 1 M NaCl and incubated at 4°C for 1 hour on a rotator. Following centrifugation at 2,500 × g for 10 minutes, the supernatant was collected as the NaCl fraction (NaCl sup). The pellet was washed again with 50 μL of PBS, and the supernatant was collected as the Wash 2 fraction. Subsequently, the pellet was resuspended in 50 μL of 100 mM NaOH and incubated at 4°C overnight on a rotator. The suspension was centrifuged at 9,100 × g (10,000 rpm) for 5 minutes, and the supernatant was neutralized with 8.0 μL of 0.6 M HCl to generate the NaOH supernatant fraction (PT fraction). The remaining pellet was subjected as pellet fraction (ppt). Protease inhibitors were added to all solutions used in the procedure.

### Protein sequence alignment

Amino acid sequence of CT83/KK-LC-1 and Gm28269 deposited on Ensembl genome browser or NCBI genome browser were downloaded. KK-LC-1 of Human (*Homo sapiens*, ENST00000371894.5, NM_00107978.4), Pika (*Ochotona princeps*, ENSOPRG00000015595), Rabbit (*Oryctolagus cuniculus*, ENSOCUG00000036103), Tree Shrew (*Tupaia belangeri*, ENSTBEG00000015209), Thirteen-lined ground squirrel (*Ictidomys tridecemlineatus*, XM_040284632), Woodchuck (*Marmota monax*, XM_046420945), Gray squirrel (*Sciurus carolinensis*, XM_047535296), Eurasian red squirrel (*Sciurus vulgaris*, ENSSVLG00005017691), Damara mole rat (*Fukomys damarensis*, ENSFDAT00000024513.1), Domestic guinea pig (*Cavia porcellus*, XM_005002715.1), Long-tailed chinchilla (*Chinchilla lanigera*, ENSCLAG00000004474), American beaver (*Castor canadensis*, XM_020152280.1), Ords kangaroo rat (*Dipodomys ordii*, XM_013019741.1), Banner-tailed kangaroo rat (*Dipodomys spectabilis*, XM_042665826), Pacific pocket mouse (*Perognathus longimembris pacificus*, XM_048335325.1), Chinese hamster (*Cricetulus griseus*, XM_007649237.1), Golden hamster (*Mesocricetus auratus*, XM040743875.1), Desert hamster (*Phodopus roborovskii*, XM_051178887.1) and Gm28269 from Mouse (*Mus musculus*, ENSMUST00000189511.1), Algerian mouse (*Mus spretus*, MGP_SPRETEiJ_T0094911.1), Steppe mouse *(Mus spicilegus*, ENSMSIT00000000470.1), Ryukyu mouse (*Mus caroli*, MGP_CAROLIEiJ_T0091163.1), Shrew mouse (*Mus pahari*, MGP_PahariEiJ_T0088803.1), Rat (*Rattus norvegicus*, EDM10792.1) were used. Multiple sequence alignment was performed with CLUSTALW. Alignment and phylogenetic reconstructions were performed using the function “build” of ETE3 3.1.2 [38] as implemented on the GenomeNet (https://www.genome.jp/tools/ete/) and plotted with JalView (https://www.jalview.org/) [39]. The tree was constructed using FastTree v2.1.8 with default parameters [40]. Protein sequences were analyzed by SMART with default parameters (http://smart.embl-heidelberg.de/) [41, 42].

### RT-PCR

Total RNA was isolated from snap-frozen tissues with ISOGEN (FUJIFILM Wako Pure Chemical) and subjected to first strand cDNA synthesis with SuperScript III cDNA synthesis kit (Thermo Fishe Scientific). Ex Taq (Takara Bio) was used for PCR reaction. PCR primers for RT-PCR was as follows, 5’-ACCAGACTCTGTTTGGTCTG-3’ and 5’-ATGTTGCCACAAACAACTGC-3’ for *Gm28269/Kk-lc-1 allele*, 5’-TGTCCGTCGTGGATCTGAC-3’ and 5’- TTACTCCTTGGAGGCCATGT -3’ for *Gapdh*, 5’-GAGAAACACCGGCGAGATG-3’ and 5’-GTTCCAGTTCCACCAGCTTGTT-3’ for synthetic *CT83*/*KK-LC-1* (*synKK-LC-1*) and 5’- GCTGATCCTGATCATCAGCC-3’ and 5’- GCTGTTGGTCTTGACGCTG-3’ for synthetic *Gm28269/Kk-lc-1* (*synKk-lc-1*).

### Immunoblotting

Proteins were separated by SDS-polyacrylamide gel electrophoresis and transferred to Immobilon PVDF membranes (Merck). After blocking membrane with PVDF Blocking Reagents for Can Get Signal (TOYOBO) or 5% Skim-milk, membrane was probed with primary antibodies overnight at 4 ℃, and subsequentially reacted with secondary antibody conjugated with horseradish peroxidase (Jackson ImmunoResearch). Anti-Gm28269/Kk-lc-1 antibody (1:5,000) was generated from rabbit serum which is immunized recombinant Gm28269/Kk-lc-1 polypeptide (VKIIKKPTRKPDKDSDSDGNN). Anti-KK-LC-1 monoclonal antibodies (1:1,000) were purchased (ac219971, Abcam) or generated in-house [43]. Antibodies used were the following: ACTRT2 1:2,000 (16992-1-AP, Proteintech), Basigin 1:200 (sc-9757, Santa Cruz), ADAM3 1:200 (sc-365288, Santa Cruz), Actin 1:1,000 (MAB1501, Millipore), ꞵ-Tubulin 1:20,000 (66240-1-Ig, Proteintech) and histone H3 1:1,000 (MABI0301, MAB institute Inc.). The signals of Anti-KK-LC-1 were detected using ECL prime Western Blotting Detection Reagent (Cytiva). Other signals were detected using 1:1 mixture of lab-maid ECL A (100mM Tris-HCl pH 8.5, 0.4mM p-Coumaric acid, 5mM Luminol) and ECL B (100mM Tris-HCl pH 8.5, 0.04% H_2_O_2_).

### *In situ* Hybridization chain reaction (HCR) with mouse and human testis

Formalin-fixed paraffin embedded (FFPE) mouse and human testis sections were used for modified *in situ* HCR [44]. Testes were removed from euthanized mice and fixed overnight in 10% formalin neutral buffer solution at 4°C. Normal human testis tissue slides (#TEN601) were purchased from TissueArray.Com LLC. The split probes for *in situ* HCR were designed with the full length of *Gm28269/Kk-lc-1* and *CT83/KK-LC-1* mRNAs, including UTR and CDS (Supplemental table 1). FFPE sections were deparaffinized with xylene and rehydrated to distilled water by graded ethanol series. They were permeabilized with methanol for 10 minutes at room temperature, washed by PBS with 0.1% Tween 20 (PBST) twice. After washing, the sections ware prehybridized for 10 minutes at 37°C with a hybridization buffer solution [1×saline sodium citrate solution (SSC) (NIPPON GENE), 10% sodium dextran sulfate 500,000 (FUJIFILM Wako Pure Chemical), 0.1% Tween 20, 50 μg/mL heparin (Nacalai tesque) and 1×Denhardt’s Solution (Nacalai tesque)]. Then, the sections were incubated overnight at 37°C, with a hybridization buffer solution containing a mixture of 20 nM split probes. After the hybridization, the slides were washed three times for 10 minutes in 0.5×SSC with 0.1% Tween 20 (0.5 × SSCT). HCR amplifications were performed using biotin-labeled hairpin DNA (IPL-BI-A161CM; Nepa Gene). The sections were incubated with amplification buffer solution (8 × SSC, 100 mM MgCl_2,_ 10% dextran sulfate 500,000 and 0.2% Triton X-100) containing the hairpin DNA pairs for 2 hours at 25℃. Then, the slides were washed with PBST three times for 10 minutes at 37 ℃ and rinsed with PBS for 5minutes room temperature. For brightfield observation with biotin-labelled hairpin DNA, the sections were reacted with Streptavidin-PolyHRP20 (Stereospecific Detection Technologies) at room temperature for 30 minutes and staining was performed with ImmPACT DAB (Vector Laboratories). They were then dehydrated and permeabilized, sealed and observed. The staining sections were imaged under an optical microscope equipped with Visualix V500FL (Visualix). In the detection of each target gene, the negative control followed the same procedure but omitted the split probe on one side.

### Staining sperm mitochondria and immunostaining of testis section

Spermatozoa were collected from cauda epididymis and dispersed in HTF medium (FUJIFILM Irvine Scientific) for 10 minutes. Incubated spermatozoa were spotted onto a glass slide and air dried. Mitochondria and DNA were stained for 15 minutes with MitoBright LT Red (DOJINDO LABORATORIES) and Hoechst 33342 (DOJINDO LABORATORIES), respectively. Immunostaining was performed using testis tissue sections prepared as described above. In the tissue sections, antigen retrieval was performed by treating the sections with ImmunoSaver (FUJIFILM Wako Pure Chemical), followed by heating for 45 minutes at 98 ℃. Then the sections were blocked with Blocking One Histo (Nacalai Tesque) for 10 minutes at room temperature. Then the tissue sections were incubated with anti-Gm28269/Kk-lc-1 (1:1000) or anti-KK-LC-1 (1:1000) antibodies for overnight at 4℃, followed by incubation for 1 hour at room temperature with donkey anti-rabbit IgG Alexa Fluor 594 (1:500, #A32754; Thermo Fisher Scientific) or EnVision+ Dual Link System-HRP (Dako) as secondary antibodies. For fluorescence detection, DAPI and Lectin-PNA-Alexa 488 (Thermo Fisher Scientific) were also added to the secondary antibody reaction solution. Autofluorescence was quenched using the Vector True VIEW autofluorescence quenching kit (Vector Laboratories). Staining was performed with Liquid DAB+ Substrate Chromogen System (Dako) according to the manufacturer’s instructions. Images were captured using the A1R HD25 confocal laser scanning microscope (Nikon) or optical microscope described above.

### Generation of *Gm28269*/*Kk-lc-1* knock-out mice

For generating *Gm28269/Kk-lc-1* KO mice, Cas9 protein (Integrated DNA Technologies) and two gRNA (Thermo Fisher Scientific) mixtures were electroporated with fertilized embryo [45] constructed by *in vitro* fertilization of C57BL/6J sperm and C57BL/6J oocytes. The gRNA 1 sequence is 5’-TATGAGGTTTACGTTTATCA-3’ and gRNA 2 sequence is 5’-AGTGGTGTTCGACGTGGTCC-3’. These two gRNAs with fewer off-target sites were designed utilizing the online tool of CRISPRdirect [46]. *Gm28269*/*Kk-lc-1* KO mice were maintained on a C57BL/6J background.

### Generation of *CT83*/*KK-LC-1* and *Gm28269*/*Kk-lc-1* knock-in mice

Fertilized embryos were generated by *in vitro* fertilization with superovulated embryos obtained from *Kk-lc-1* homologous KO females and WT C57BL/6J sperm. Fertilized embryos were electroporated with Cas9 protein and gRNA with synthetic *CT83*/*KK-LC-1* (synKK-LC-1) or synthetic *Gm28269/Kk-lc-1* (synKk-lc-1) single-stranded oligodeoxynucleotide (ssODN; Fasmac). Sequence of gRNA for ssODN knock-in (KI) is as follows 5’-CTGATGCTTGCTTTTTACCA-3’. This gRNA is designed from a sequence obtained from *Kk-lc-1* knock-out mice which harbor frameshift for *Kk-lc-1* (Supplemental Fig. 3A) by CRISPRdirect. Nucleotide sequence of synKK-LC-1 and synKk-lc-1 ssODN is described in supplemental information (Supplemental Fig. 3B and C). KI mice were maintained on a C57BL/6J background and F2 and subsequence generation were used for the experiments.

### Fertility test

For testing *in vivo* fertility, sexually matured (7-12 weeks of age) KO male mice or WT male mice were caged with female C57BL/6J mice (7-12 weeks of age) for 4 months. The number of pups in a pregnancy was calculated as litter size.

### *In vitro* fertilization (IVF)

Spermatozoa collected from cauda epididymis were incubated in HTF medium for 2 hours at 37 ℃ under 5% CO_2_ in air for sperm capacitation. Eggs were collected from superovulated C57BL/6J females (3-4 weeks of age) 17-18 hour after hCG injection. Capacitated sperm were mixed with cumulus-oocyte complex (COCs) suspended in HTF drop (final concentration: 400 cells/μL). The fertilization rate was counted as the developmental rate to the 2-cell stage approximately 20 hours after sperm addition. For zona (ZP) removal, COCs were treated with hyaluronidase (FUJIFILM Wako Pure Chemical) to remove cumulus cells, and the cumulus-free oocytes were moved to the drop containing acidic Tyrode’s solution (FUJIFILM Wako Pure Chemical) for a few seconds to remove ZP. After ZP-removal, oocytes were washed and transferred to HTF medium and incubated for 1 hour before insemination.

### Measurement of sperm motility by sperm motility and morphology analysis system (SMAS)

Epididymal sperm from WT and *Gm28269/Kk-lc-1* KO sperm (11-12 weeks of age) were dispersed in BSA-free TYH medium and incubated for 5 minutes. After 5 minutes of incubation, portions of the sperm drop were transferred to TYH drop containing BSA then incubated for 0-hour, 0.5-hour, 1 hour, 2 hours and 3 hours. At each timepoint, more than 200 sperm were collected and analyzed for motility by SMAS (DITECT). Scores for VAP (Average path velocity in μm/sec), VCL (Curvilinear velocity in μm/sec), VSL (Straight-line velocity in μm/sec), ALH (Amplitude of lateral head displacement in μm) and BCF (Beat cross frequency in Hz) were obtained at each timepoint by SMAS. For calculating hyperactivated sperm from these parameters in accordance with the criteria of Goodson, the following formulas were used to perform the calculations. SVM2 = (0.0123 × VAP) – (0.1034 × VSL) + (0.0307×VCL) + (0.0427×ALH) + (0.0175×BCF) – 3.6222. If the score of SVM2 was greater than 0, the sperm was considered hyperactivated. The sperm flagellar bending patterns were traced from 30 frames from video tapes recorded at 180 frames per second using SMAS. This process was performed using Bohboh (Bohbohsoft, Tokyo, Japan, https://documentary-ch.com/product/software/bohboh/), a software for analyzing cell motility. The amplitude of flagellar curvature at a location 5.5 µm from the connection between the sperm head and tail was calculated and plotted.

### Artificial insemination

Superovulated C57BL/6J females (8-12 weeks of age) were artificially inseminated with cauda epididymal sperm 12 hours after hCG injection. Cauda epidydimal sperm were capacitated in HTF medium for 2 hours at 37 ℃ under 5% CO_2_ in air. Capacitated sperm suspension (about 1.375-4.2 × 10^5^ sperm/5-10 μl) were injected into the uterus at a distance of approximately 1 cm from the uterotubal junction with glass microcapillary [19]. The oviducts were exercised and flushed with HTF medium to recover unfertilized oocytes or 2-cell embryos 24 hours after sperm injection.

### Reanalysis of single-cell RNA-seq data from testicular cells

The mRNA expression of CT83/KK-LC-1 and Gm28269/Kk-lc-1 in spermatogenic cells was analyzed in silico by Loupe Browser (10x Genomics) with single-cell RNA -seq dataset published previously [47].

### Artificial Intelligence (AI)

During the preparation of this manuscript, the authors used ChatGPT-5.2 (Open-AI) to refine wording, improve grammatical accuracy, and enhance overall readability. All content was subsequently reviewed and edited by the authors, who take full responsibility for the final version of the manuscript.

## Acknowledgement

We thank Drs. Shin-ichi Kashiwabara and Yoshinori Kanemori for their valuable comments on the detection of ADAM3. This work was supported by JSPS KAKENHI Grant Numbers JP22K05562 (T.Y.) and JP22H04926 (Grant-in-Aid for Transformative Research Areas-Platforms for Advanced Technologies and Research Resources “Advanced Bioimaging Support” (K.I.). This work was also supported by the ALL KITASATO PROJECT STUDY (T.F.) and the Science Research Promotion Fund from the Promotion and Mutual Aid Corporation for Private School of Japan (T.F.).

---

**Supplemental Figure 1.**
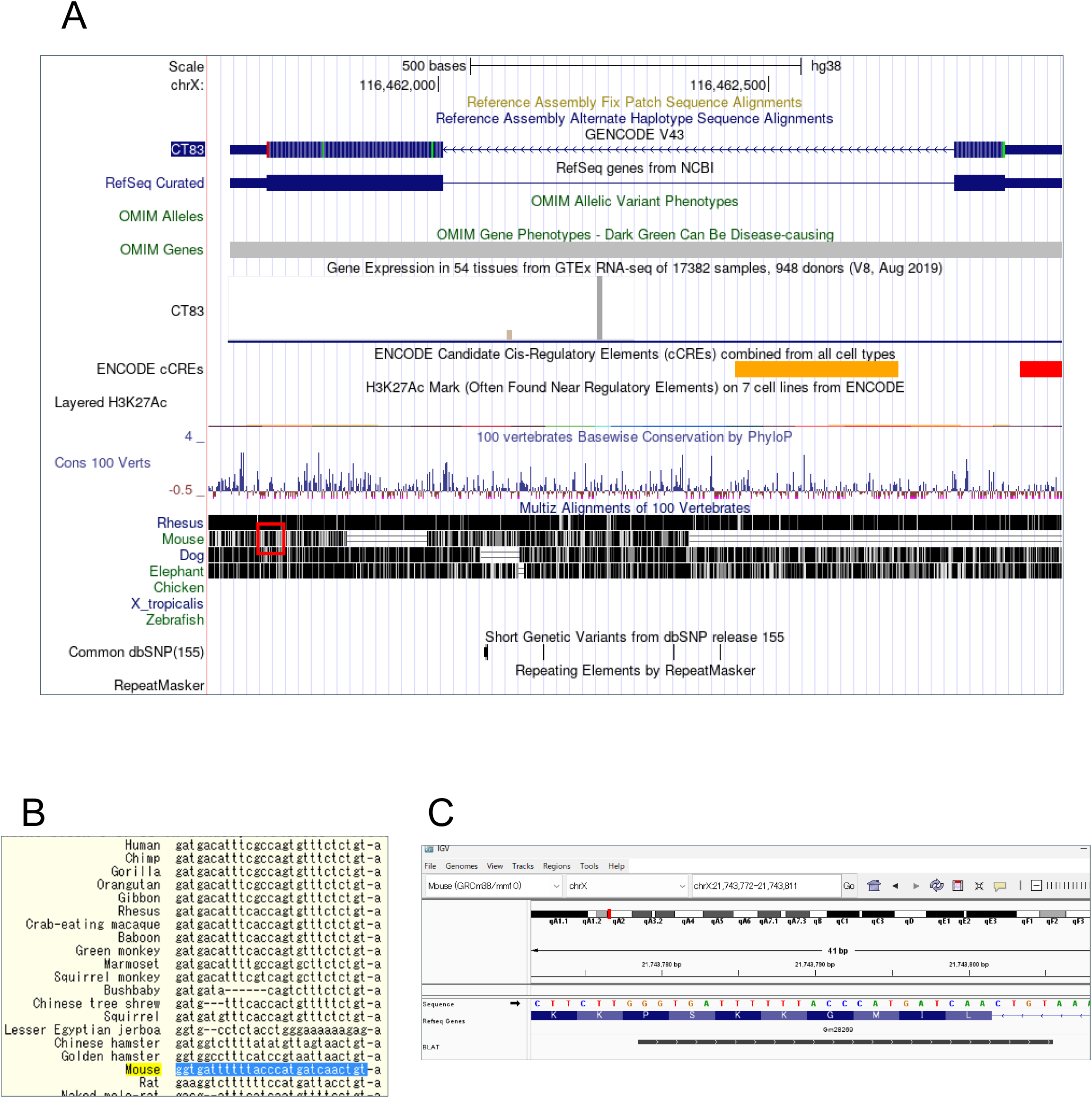
Identification of mouse homolog of *KK-LC-1*. A. A snapshot of *CT83*/*KK-LC-1* on the UCSC genome browser. In the Multiz Alignments of 100 Vertebrates track for mice, regions with high homology to the human sequence were highlighted with a red outline. B. DNA sequences highly identical to 3’ region of *CT83*/*KK-LC-1* in various species. Sequence of mouse was highlighted with blue. Note that sequences of primates were highly identical to human sequences. C. Identification of candidate genes through BLAT search in IGV. Identified sequence “GGTGATTTTTTACCCATGATCAACTGT” was BLAT searched with IGV genome browser. A candidate gene of *CT83*/*KK-LC-1* mouse homolog *Gm28269* was identified.

**Supplemental Figure 2.**
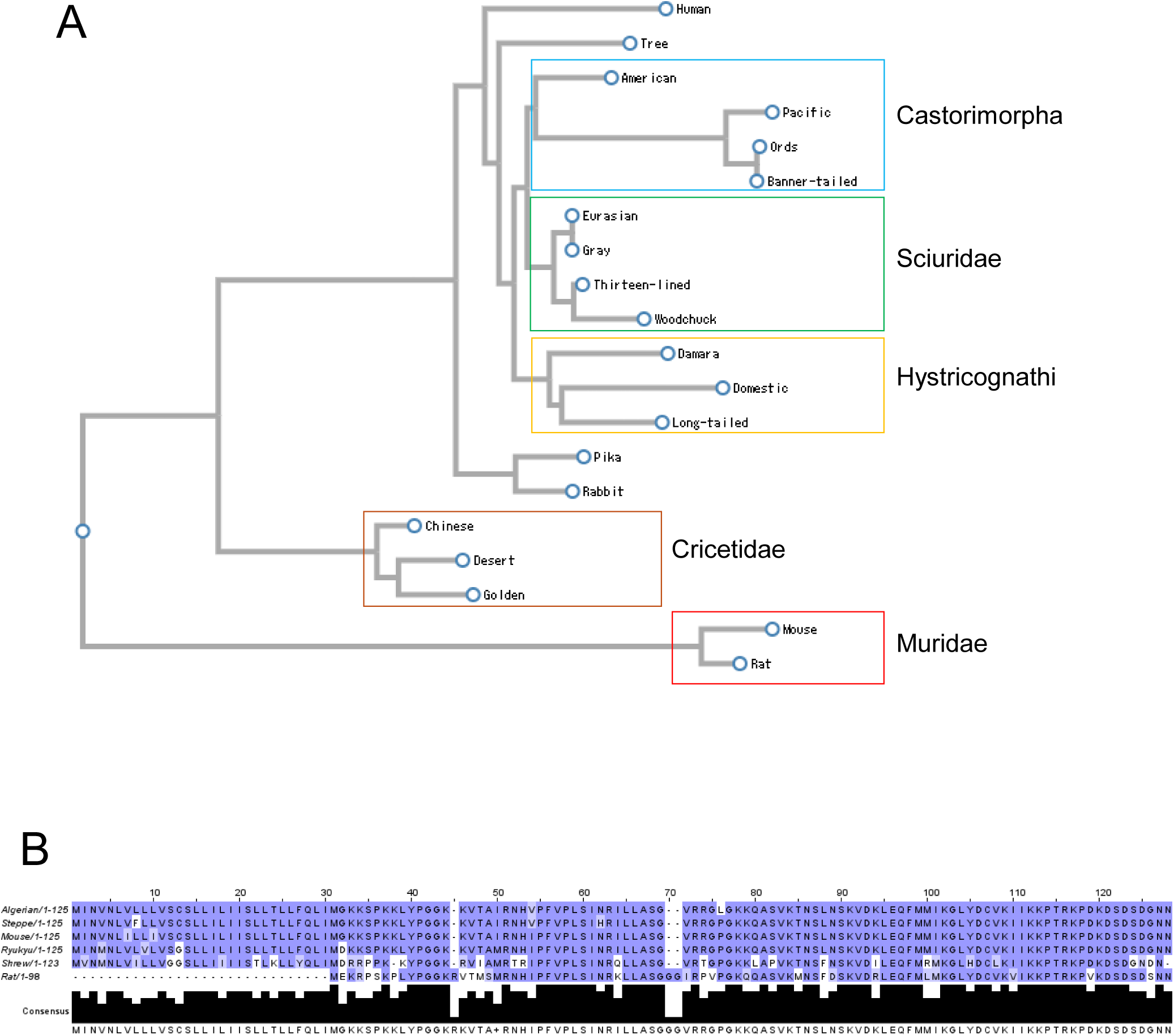
Characterization of *Gm28269*/*Kk-lc-1*. A. Amino acid phylogenic tree of *CT83/KK-LC-1* and *Gm28269/Kk-lc-1*. Among the rodent orders, the suborder Castorimorpha (blue), Sciuridae (green), Hystricognathi (yellow), Cricetidae (brown) and Muridae (red) are circled. B. Amino acid alignment of *Gm28269*/*Kk-lc-1* obtained from species from genus mus by CLUSTALW. The amino acids were color-coded according to the Blosum62 matrix. Gaps are colored white. If a residue matches the consensus sequence residue at that position it is colored dark blue. If it does not match the consensus residue but the 2 residues have a positive Blosum62 score, it is colored light blue.

**Supplemental Figure 3.**
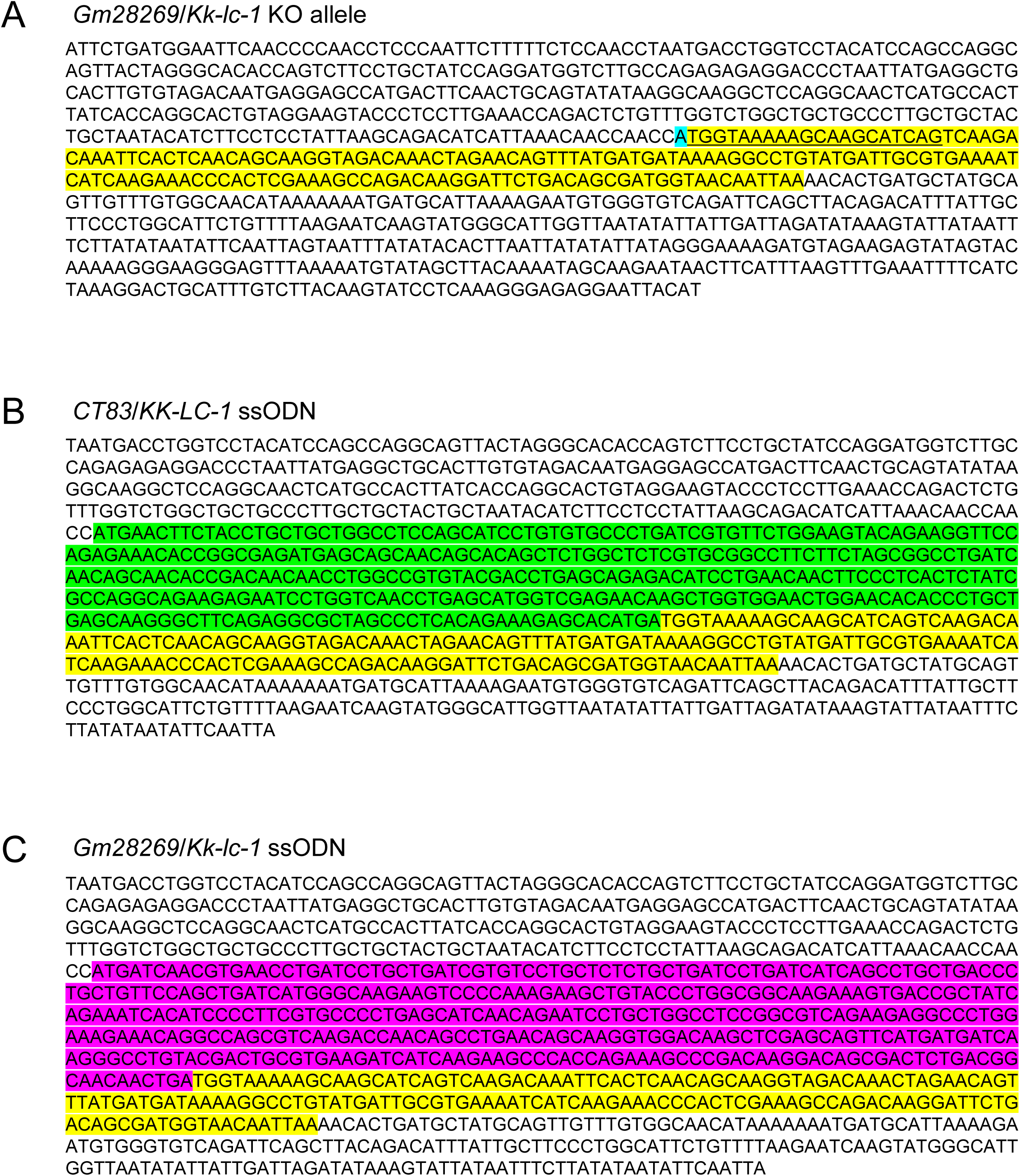
Nucleotide sequences of ssODN to produce *CT83*/*KK-KC-1* and *Gm28269*/*Kk-lc-1* KI mice. A. The nucleotide sequence of the *Kk-lc-1* locus in *Kk-lc-1* KO mice. The region highlighted in cyan represents nucleotides derived from exon 1, while the region highlighted in yellow represents nucleotides derived from exon 2. The underlined region indicates the gRNA design site for producing KI mice. B. *CT83/KK-LC-1* ssODN. The nucleotides highlighted in green represent the codon-optimized sequence of *KK-LC-1*. The nucleotides highlighted in yellow represent original exon2 sequences that remains after knockout of *Kk-lc-1*. C. *Gm28269/Kk-lc-1* ssODN. The nucleotides highlighted in magenta represent the codon-optimized sequence of *Kk-lc-1*. The nucleotides highlighted in yellow represent original exon2 sequences that remains after knockout of *Kk-lc-1*.

## Notes

### Competing Interest Statement

The authors have declared no competing interest.

